# Eye-movements as a signature of age-related differences in global planning strategies for spatial navigation

**DOI:** 10.1101/481788

**Authors:** Elisa M. Tartaglia, Celine Boucly, Guillaume Tatur, Angelo Arleo

## Abstract

The ability to efficiently find alternatives routes when faced with unexpected obstacles along our path is among the most compelling evidence of the flexibility of human behaviour. Although a plethora of plausible computations have been put forward to elucidate how the brain accomplishes efficient goal-oriented navigation, the mechanisms that guide an effective re-planning when facing obstructions are still largely undetermined. There is a fair consensus in postulating that possible alternatives routes are internally replayed sampling from past experiences, however, there is currently no account of the criterion according to which those memories are replayed. Here, we posit that paths, which are expected to be more rewarding are replayed more often and that eye movements are the explicit manifestation of this re-planning strategy. In other words, the visual sampling statistics reflects the retrieval of available routes on a mental representation of the environment.

To test our hypothesis, we measured the ability of both young and old human subjects to solve a virtual version of the Tolman maze, while we recorded their eye movements. We used reinforcement learning (RL) to corroborate that eye movements statistics was crucially subtending the decision making process involved in re-planning and that the incorporation of this additional information to the algorithm was necessary to reproduce the behavioral performance of both screened populations.

## Introduction

The ability to efficiently find alternatives routes when faced with unexpected obstacles along our path is among the most compelling evidence of the flexibility of human behaviour. If, for example, our usual way to the bakery is blocked by some construction work, we would still be able to reach our destination by making -hopefully-a small detour.

Although a plethora of plausible computations have been put forward to elucidate how the brain accomplishes efficient goal-oriented navigation (for a review see Madl et al. 2015; Chersi and Burgess 2015), the mechanisms that guide an effective re-planning when facing obstructions are still largely undetermined.

There is a fair consensus in postulating that possible alternatives routes are internally replayed sampling from past experiences via network interactions involving hippocampus and prefrontal cortex (Spiers and Gilbert 2015). However, there is currently no account of the criterion according to which those memories are replayed. Here, we posit that paths which are expected to be more rewarding are replayed more often and that eye movements are the explicit manifestation of this re-planning strategy. In other words, the visual sampling statistics reflects the retrieval of available routes on a mental representation of the environment (i.e. on a cognitive map).

To test our hypothesis, we measured the ability of human subjects to solve a virtual version of the Tolman maze, while we recorded their eye movements. In his original study, Tolman found that rodents were capable to take detours and to select the shortest path to food pellets. Consequently, he put forward that an internal representation of the physical space is a necessary condition to be able to re-plan one’s way in a modified environment (Tolman 1948). In the light of these results, we expected young adults not only to be able to successfully re-plan their way through the maze, but also to be able to sketch a faithful representation of the maze, as further evidence of a cognitive map formation (one that cannot be accounted for in non-human subjects). According to our hypothesis, then, the eye movements statistics collected when subjects are required to find alternative routes, should reflect planning over the cognitive map; if such map is correct, the correct alternative route will be chosen.

A much more challenging instance to test our hypothesis upon concerns older adults performance. Aside from general spatial navigation deficits (Barrash 1994; Wilkniss et al. 1997; Burns 1999; Newman and Kaszniak 2000; Moffat and Resnick 2002; Driscoll et al. 2005; Head and Isom 2010; Jansen et al. 2010; Lester et al. 2017), aging has been specifically linked to a delay in cognitive map formation as well as to an impediment in using it (Iaria et al. 2009). Moreover, elderly have been shown to be defective in allocentric navigation (Moffat et al. 2006; Antonova et al. 2009; Rodgers et al. 2012; Bohbot et al. 2012; Gazova et al. 2013), which would suggest that they rely more on a stimulus-response association type of navigation strategy, rather than on planning. According to our hypothesis, instead, older adults, as their younger counterpart, are capable to do mental planning; likewise, their eye movements reflect planning over a cognitive map. However, we conjectured, older adults re-plan their alternative route over a degraded cognitive map; this might explain why older subjects have been found to be impeded in solving a detour task and/or in taking shortcuts (Harris and Wolbers 2014).

To corroborate that eye movements statistics was crucially subtending the decision making process involved in re-planning, we used reinforcement learning (RL) models. Although a phenomenological model, both neuroimaging and neurophysiological studies have provided strong indication that specific anatomically identified brain regions are indeed performing RL based computations (Schultz et al. 1997; Berns et al. 2001; O’Doherty 2004; Pagnoni et al. 2002). Moreover, the use of RL to fit behavioural data of (young) observers performing a navigation task has provided a clear indication of the strategies at use. In particular, contrasting model-based and model-free RL algorithms have led to identify whether navigation strategies involve or not the ability of planning future path selection (Daw et al. 2005; Simon and Daw 2011; Keramati et al. 2011; Gershman et al. 2014; Tartaglia et al. 2017).

Here, we show that the incorporation of EM statistics as additional information to RL algorithms was necessary to reproduce the behavioral performance of both screened populations. In the classical RL theory, the *agent* (e.g. the observer navigating through the maze) learns from reinforcement the future value of the expected reward at any given *state* of the environment (e.g. a location in the maze). The value of a state is updated only if the agent has visited it (Sutton and Barto 1998). Here, we made a much stronger assumption: the states that observers looked at are used for planning and consequently updated even if they have not been visited. The rationale behind this assumption goes as follow: often, in a natural setting, visual exploration of the environment alone provides enough information about which direction should be taken next. Imagine you are at a crossroad, looking to the right you realize the street is dead-ended, hence you plan to turn left instead. In classical RL, the state values on the right would not get updated since the agent did not visit them. However, we argued, a more efficient and biologically plausible algorithm should be able to assign (in this case negative) values to the right path.

More into details, using the recorded data, we fit our RL algorithm to the behavioural choices (i.e. the discretised sequence of maze locations visited by each observer), as well as to the number of fixations performed during the test trials by both young and old observers. In particular, as a first step, we fed to the model (with two free parameters) the sequence of states visited during the exploration phase by each individual subject. This method provided individual (and more realistic) initial conditions to be used, in a second step, in the test trials, during which we compared observers’ performance to the model performance. Importantly, the model was enriched with a crucial information: the statistics of eye movements made by each individual subject when planning their next move towards the goal. As a final step, we determined the region of parameters which yielded the closest performance to the behavioral data.

We seek to provide, for the first time, a plausible mechanistic account of the process of planning alternative routes when facing unexpected obstacles. Furthermore, we expect these results to lead to a quantification of the extent to which such process differ from young to older observers.

## Methods

### Participants and set-up

We tested *n* = 39 participants. Age and gender of the sample are reported in the following table.

| Gender | Young (< 45 years) | Old (>65 years) |
| --- | --- | --- |
| Female | 9 | 10 |
| Male | 10 | 10 |

The mean age for young and old observers was, respectively, 27.3*y* and 78.5*y*. Experiments were run on a simulation module developed in Unity 3D, which allowed to create the virtual maze and to record subjects trajectories. A view from above of the maze, together with its dimensions is shown in Fig 1A. A chest at the end of the maze was used as Goal (Fig 1B). Any visual cue within the environment, which could help subjects to reach the Goal faster, was avoided.

**Figure 1:**
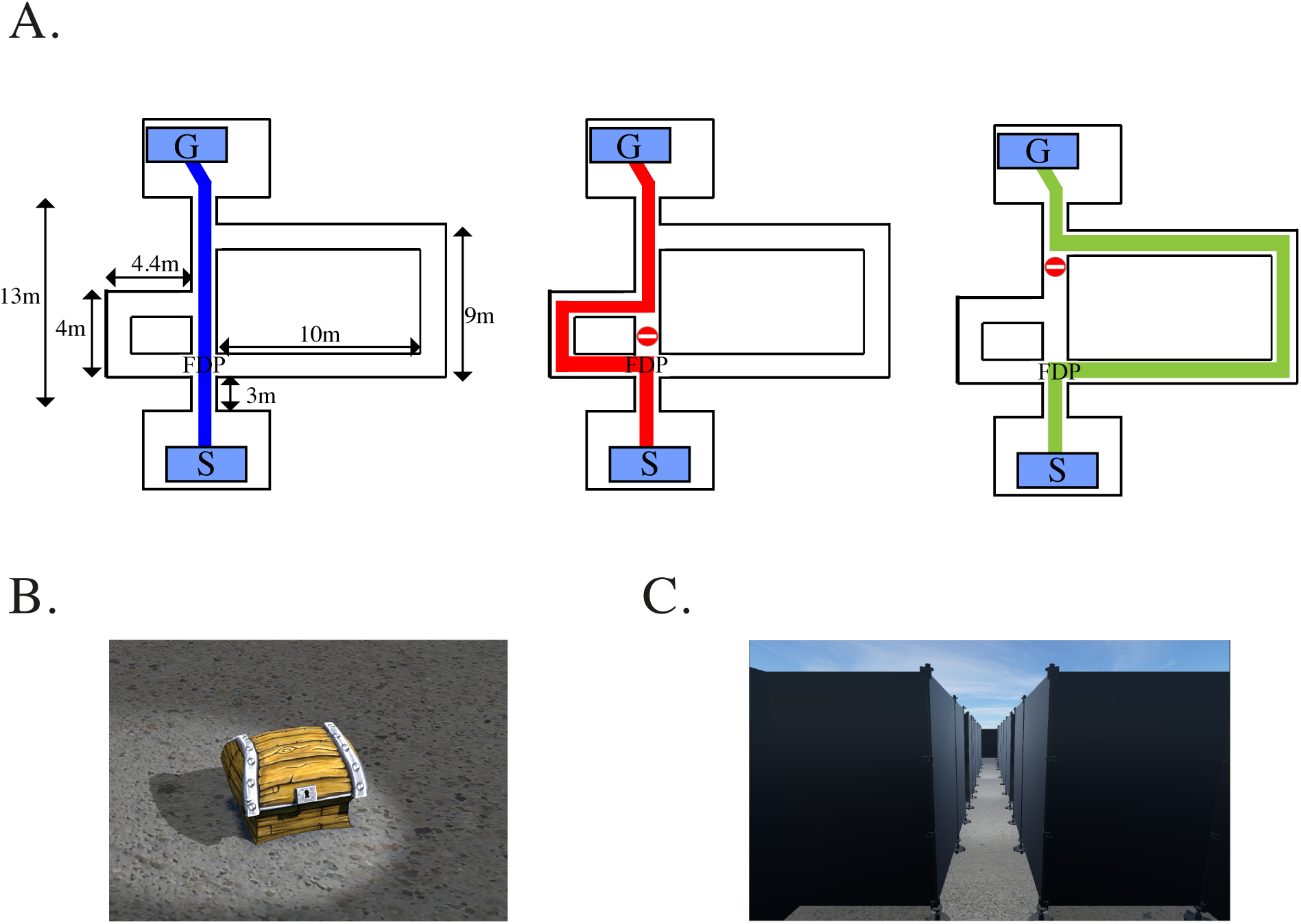
A. Shortest paths from Start (labelled as *S*) to Goal (labelled as *G*) in the three conditions of the test trials: from left to right, no block (0), block A, block B respectively. *FDP* labels the first decision point in the maze. The actual dimensions of the maze are also indicated. **B.** A picture of the chest located in *G*. **C.** Simulator view of the maze from the starting point *S* in the subjects’ prospective.

Three main paths allowed to reach the chest: a central corridor, a left and a right path, rejoining the central corridor at different locations (exit points). The central corridor provided the shortest path to the Goal, while the right one provided the longest (Fig 1A).

Exit points of both right and left paths were hidden by virtual doors, to prevent participants to get a glimpse of the maze structure from the very starting point; virtual doors opened only when subjects were in their close proximity and were not identifiable otherwise.

Subjects were generally allowed to take any path, from any direction, within the maze (except when specified otherwise).

The virtual maze was presented on a projection screen (0.9*m* ∗ 1.8*m*). Subjects were sitting at a distance of 1.5*m* from the screen and moved within the environment –from a first person perspective-using a joystick. Turning rate and walking speed were kept constant i.e. all participants navigated within the environment at the same speed. This allowed to minimize differences in performance that might arise from age-related difficulties in using the joystick. We synchronized the virtual walking with footsteps sound, to render the virtual experience more realistic.

Binocular eye movements were recorded throughout the experiment with a head-mounted eye tracking system (Eyelink *II*, SR Research). The calibration of the eye tracking was done before the experiment and was regularly checked upon during the whole experimental session.

## Procedure

Before giving their consent, subjects were informed that they were going to participate, for about 1.5 hours, in a virtual navigation study, in which their eye movements would have been recorded. The experiment was divided into three main phases: *training*, *exploration* and *testing*.

During training, instructions were verbally given and assistance with joystick manipulation and task understanding was constantly provided. During both exploration and testing phase, instructions appeared on the screen to minimize any experimenter bias.

### Training phase

To reduce motor skills differences, we asked subjects to practice the use of the joystick while navigating in a virtual environment -which was different from the Tolman maze. Several repetitions of this training task were run, depending on how good the experimenter judged the individual ability to handle the joystick. Once subjects were judged at ease with the set-up, they were asked to practice a *pointing task* in yet another rendering of the virtual environment. In this case navigation was guided, i.e. subjects were asked to follow a given trajectory until a compass appeared on the screen. The task consisted in using the joystick to point the compass arrow towards a previously visited location, which was out of sight from the current position. The rationale of this phase was to get the subjects acquainted to the use of the compass in the virtual environment, before the proper pointing task in the Tolman maze was carried out. Being able to point to the correct direction provided an indication of the observers ability to locate themselves in the environment, as well as an indirect measure of their ability to form an internal representation of the environment. Eye movements were not recorded during the training phase.

### Exploration phase

Subjects were first instructed to freely navigate in the virtual Tolman maze, of whom they ignored the structure; we informed them that a chest was located somewhere in the maze, although they were not explicitly instructed to look for it, or to memorize its location. Subjects were also aware that the chest acted like a portal: once reached it, they would be instantaneously brought back to the maze starting point. In other words, the chest allowed to resume navigation from a fixed reference point, providing a useful direction cue. This first stage of exploration lasted six minutes.

Next, we instructed subjects to follow popping out arrows guiding them from the starting state to the Goal trough the three main maze paths, i.e the left, the central and the right path. This “guided exploration” phase lasted not more than two minutes and was designed to ensure that all subject went through each of the three paths at least once before the test phase.

A second session of free exploration, identical to the first one, followed.

The whole exploration phase lasted twelve minutes (such duration was determined based on a previous pilot study). Eye movements were recorded all along.

### Testing phase

We used three different kind of post-exploration tests to evaluate subjects performance, i.e. their ability to grasp the Tolman maze structure. Each test phase will be described in details in the following section.

## Performance measurements

### Detour task

Participants were instructed to reach the chest from the starting point by traveling the shortest possible distance. The structure of the maze as well as the chest location were left unchanged with respect to the exploration phase. Observers performed asequence of 7 trials, which we labelled as: 0 *-A-B-A-B-A-B*. In condition 0, all paths were available, hence the central corridor provided the shortest path to the chest; in condition *A*, a portion of the central corridor was blocked, therefore the left path became the shortest; in condition *B*, a portion of both the central corridor and the left path were blocked and the right path became the shortest way to the Goal (Fig 1A). Eye movements were recorded during the whole session.

To quantify performance we measured for each subject *i*, each trial repetition *j* (*j* = 1, 2, 3) and each trial type *k* (*k* = 0*, A, B*), the length of the distance travelled from Start to Goal, *D*_*ijk*_, to which we subtracted the shortest path length *min*(*D*_*k*_), i.e. the length of the shortest possible path leading from Start to Goal:

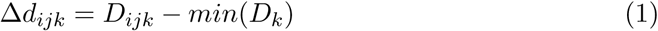

Given the above definition, Δ*d*_*ijk*_ *∈* [0*, ∞*[; performance was optimal when Δ*d*_*ijk*_ = 0.

### Sketched map task

After a quick break, participants were instructed to draw on an electronic tablet a bird’s eye view of the maze. They were asked to label the hallways, the obstacles they encountered and the chest location. They could take as long as they needed to complete the task. Eye movements were not recorded.

To quantify performance we designed a grading system and asked four unbiased raters to grade the subjects’ maps according to several criteria (see Table in Appendix and Billinghurst and Weghorst 1995). A maximum of 20 points was assigned if the maze was correctly drawn. Negative points were assigned when subjects drawn extra-elements, i.e. hallways or intersections that did not exist (see Fig. 4C).

**Figure 4:**
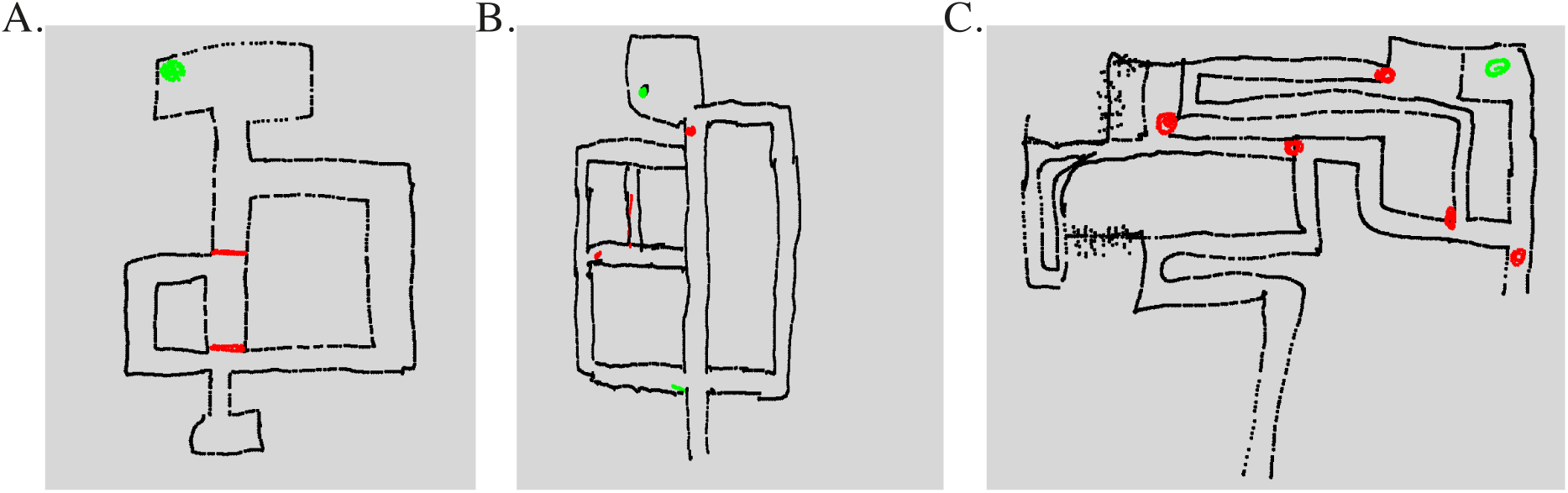
Behavioral results - Sketched map task. One young (**A.**) and two old subjects (**B., C.**) rendering of the Tolman maze. Drawing were done right after the detour task. Observers were instructed to label in green the Goal location and in red the blocks. Average grades were, from left to right, 20, 14.5 and −6.

### Pointing task

After sketching the map, participants were presented with the virtual Tolman maze again and were asked to point toward the chest in four different locations along the left and the right path (Fig. 3). We measured performance by calculating the angular deviation of each subject choice from the actual Goal location, i.e. the error subjects made in pointing towards the chest from their current position.

**Figure 3:**
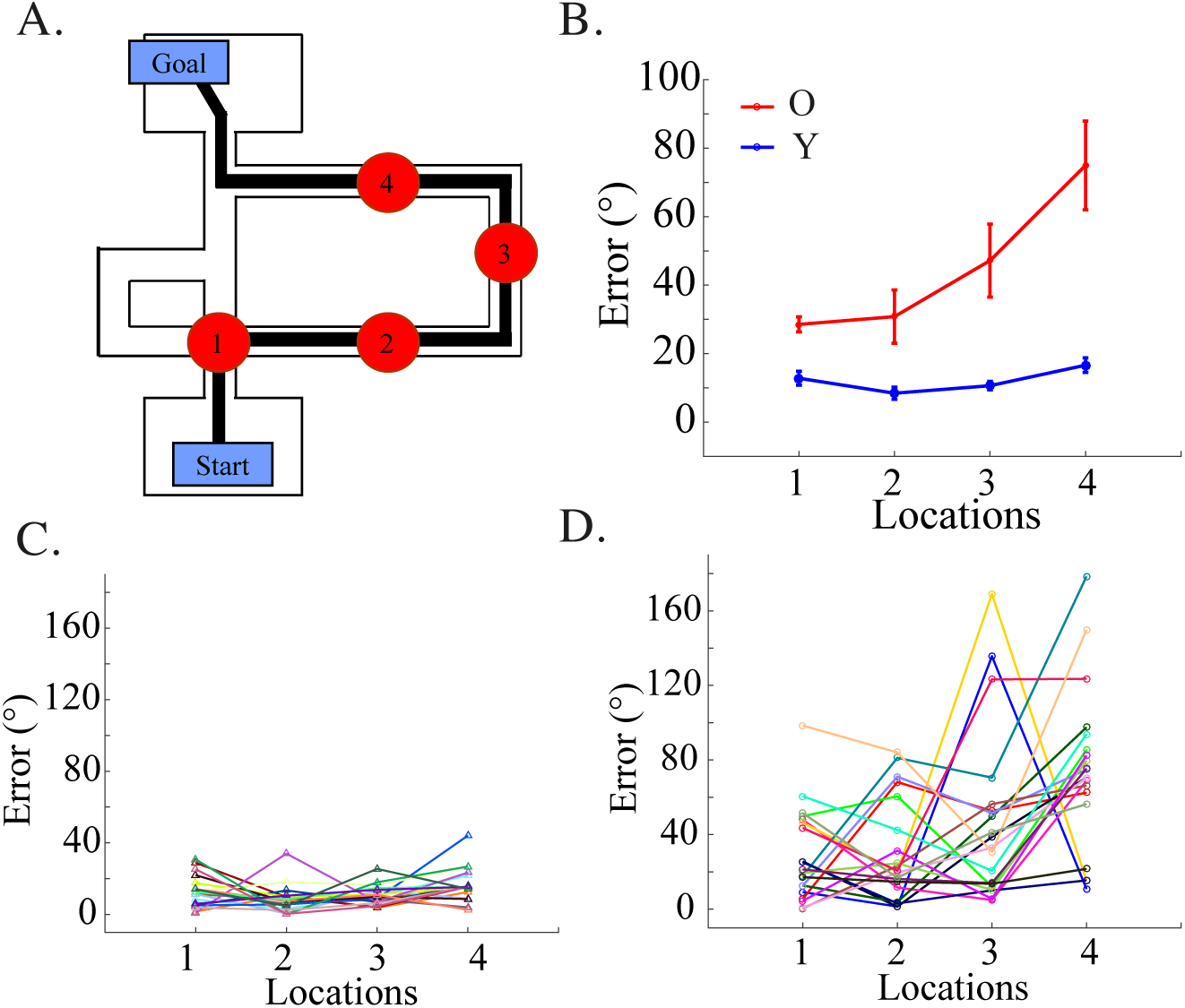
Behavioral results - Pointing task. **A.** The red circles label the four locations along the right path in which observers were required to stop and point to the Goal position. Average across subjects of the deviation from the Goal position measured in each of the four locations in A. While the error in the young population tends to remain constant (and very small), it increase with the number of changes in direction for the old population. Results are significant in any of the tested location (location 1, *p* = 0.04; location 2, *p* = 0.001; location 3, *p* = 0.0005; location 4, *p <* 10^*-*5^; Wilcoxon rank sum test). Following the same procedure, we also tested four locations along the left path and found similar results. Same as in B., but for individual young observers. **D.** Same as in C., but for individual old observers.

## Eye-movements analysis

Eye movements were recorded binocularly at a rate of 250*Hz* during both the exploratory and the testing phase. The EyeLink *II* system allowed to detect saccades and blinks by setting threshold parameters for saccade’s velocity (30°*/s*), acceleration (9000°*/s*^2^) and motion (0.15°). Periods between saccades in which blinks were absent, were identified as fixations. A fixation was quantified in terms of *gaze direction*, i.e. the angle between the centre of the virtual camera (which coincides with the position of the subject in the virtual space) and the 2D fixation point on the screen, whose coordinates were directly provided by the eye tracking device. From the point of view of a subject moving in the virtual space, the horizontal field of view spanned approximately 106°.

For the analysis, we focused on the subset of gaze directions recorded while subjects navigated from the maze starting point to the first decision point (*FDP*, see Fig. 1A), i.e. the portion of space in which subjects were expected to make a decision about which of the three paths to choose. This portion of the maze, which we referred to as *region of interest*, spanned 5.5*m*. We analyzed, separately, gaze directions recorded during the exploration phase and those recorded during each test trial, pooling together gaze directions for each population and test trials of the same kind.

We were interested in testing two hypothesis concerning eye movements dynamics: first, as a manifestation of learning, the pattern of gaze directions changed significantly when comparing the exploration phase to the test trials; second, as a manifestation of a different decision making dynamics, the pattern of gaze directions within test trials changed significantly when comparing younger to older adults. To test our hypothesis, we partitioned the region of interest in several subregions, for each of which we built a gaze direction distribution polling together the gaze direction data of the whole population. The number of subregions was determined by a sliding window of size 0.5*m* and step 0.05*m*. Such a representation allowed us to assess and to compare the evolution in time, from start to *FDP*, of the gaze direction distribution of each population.

To be able to incorporate eye movements during re-planning of an alternative route in the model, we analysed eye movements statistics for each individual observer during test trials. In particular, we counted the proportion of fixations made straight ahead towards the central corridor, towards the right and towards the left arm while the observer moved from start to *FDP*. Note that we discarded the number of times in which observer looked behind towards the starting point simply because none of the observers went back to it before reaching the *FDP*. Subsequently, we fed the proportion of eye movements to the model.

## Model

We used hierarchical reinforcement learning (HRL) to model subjects’performance in the Tolman maze task. HRL, unlike standard RL algorithms, allows the learning agent to take actions extended in space and to keep track of the distance travelled while executing them. We referred to this class of actions as *options*. Formally, each option is labelled by its initial and terminal state, as well as by its length *z* (Fig.6A).

In a given state *s*, the agent chooses an option *o* according to the following probability rule (*soft-max policy*):

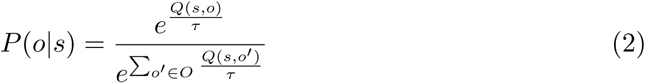

where *Q*(*s, o*) is the value associated to option *o* while in *s* and *τ* is a “noise” parameter. The normalisation takes into account all available options *o’* in state *s*. If *τ* → 0, the option with the highest probability to be selected is the one associated with the largest *Q* value. In this case, the agent is said to be *greedy*. To the contrary, the larger the *τ* the more the agent is *explorative* and is willing to choose options whose *Q* value is not necessarily the largest. In the limit of *τ* → *∞*, all available options in *s* become equiprobable.

Once the option *o* has been chosen, the *Q* value associated to the option initial state *s* is updated according to:

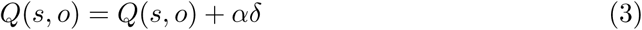

where *α* controls the learning rate and *δ* adjust the previous estimation of the reward given the reward *r* collected when executing *o* from *s*:

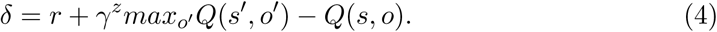

As in standard *Q*-learning algorithms, *γ* is the discount factor, which weights the rewards according to how far from *s* they will be obtained (0 ≤ *γ* ≤ 1). Unlike standard *Q*-learning, here *γ* is affected by the option length *z*, i.e. by the number of steps elapsed since *o* was chosen, so that longer options are discounted more.

The execution of option *o* brings the agent to the option terminal state *s’*. The task of the agent is to maximise the total reward across the environment through the update of *Q* values estimation. If the algorithm has converged, choosing the option associated with the largest *Q* value at each *s*, ensures the highest total reward.

In the model-free version of HRL (MF-HRL), *Q*(*s, o*) gets updated online, i.e. only after the agent visits *s* and chooses *o*. In the model-based version of HRL (DQ-HRL), *Q* values are updated also offline, through *planning*. Planning works in the following way: upon visiting *s*, the transition *s* → *o* → *s’* and its associated reward *r* are stored in a matrix *M* to build up a “model” of the environment. Before choosing the next option, the agent uses *M* to randomly sample *n*_*pl*_ states and options visited in the past and update their *Q* values offline. This planning procedure provides faster convergence to the optimal *Q* values.

In the following, we used DQ-HRL to reproduce the behavioural data. The model has four free parameters, *τ, α, γ, n*_*pl*_; we fixed *γ* = 0.9 for both young and old subjects; we assumed the remaining three parameters to differ between the two populations, but not within individual subjects.

## Results

### Behavioral Results

As expected, we found that young observers significantly outperformed old ones in the detour task. The histogram of Δ*d*_*ijk*_ shows that the whole young population performed optimally, or nearly optimally, in all trials, i.e. choosing almost always the shortest possible path to the Goal (Fig. 2A.). A much higher variability and sub-optimality was observed for older subjects, with some trajectories deviating up to 250*m* from the shortest path length (Fig. 2B.). The difference in performance between the two populations appears even more pronounced when plotting 〈Δ*d*〉_*i*_ averaged across subjects, with young subjects performing significantly better than old ones in all but one trial (Fig. 2C.,F.). Averaging further across repetitions of the same trial type (Fig. 2D.,G.), or across all trials confounded (Fig. 2E.,H.) confirmed these results. Furthermore, we found that young observers were significantly more accurate in pointing towards the Goal location in all tested sites (Fig. 3). Old subjects appear to be less and less able to locate the Goal at each change of direction in the maze (Fig. 3B). Analogous results were found when testing pointing performance across four locations spread on the left arm, or when pointing towards the starting point rather than the Goal (results not shown).

**Figure 2:**
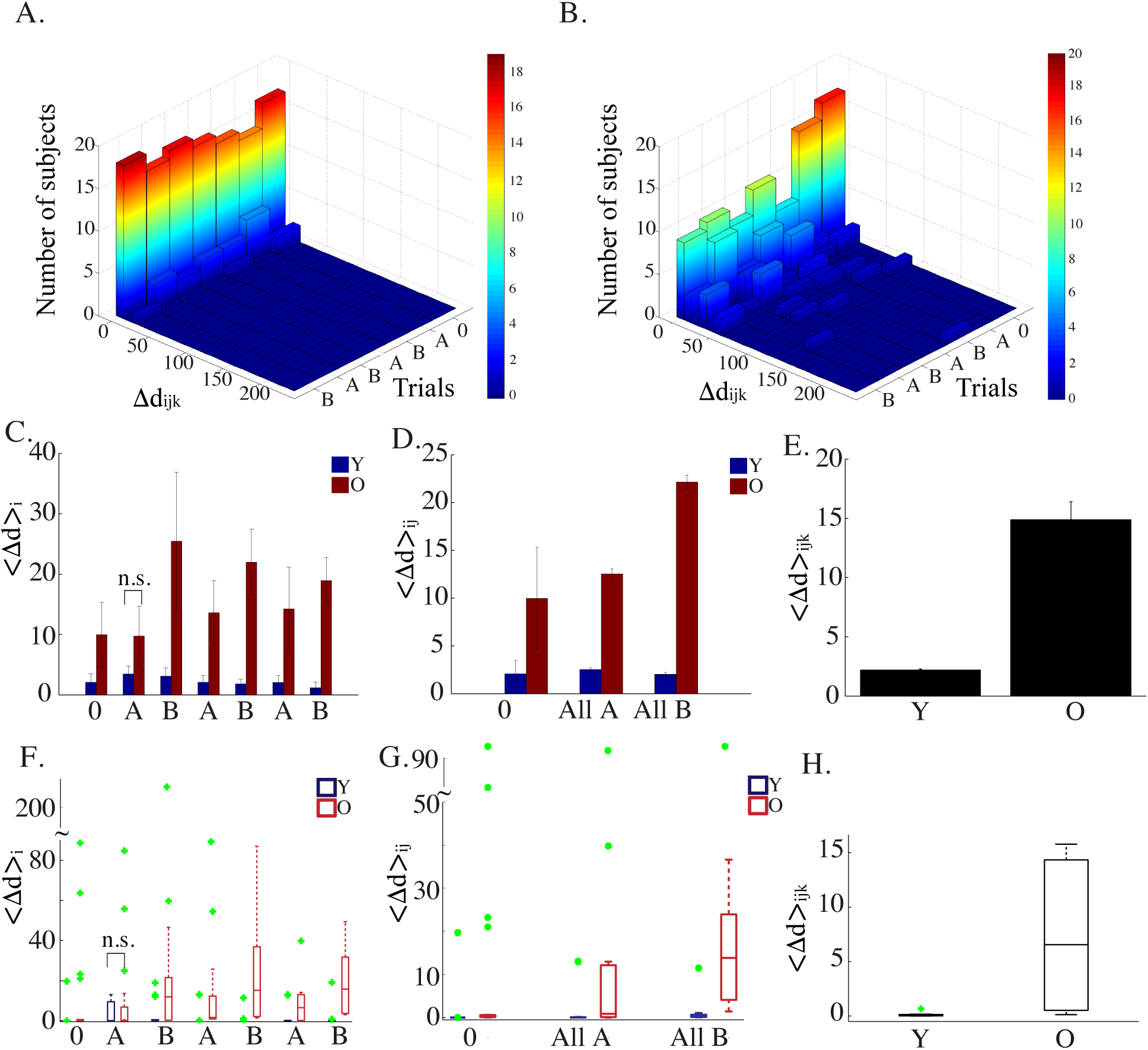
Behavioral results - Detour task. Histogram of the distance travelled from Start to Goal across the 7 trials (0-A-B-A-B-A-B) for the young (**A.**) and the old population (**B.**) An optimal performance implies Δ*d*_*i,j,k*_ = 0 (see text for more details). Travelled distance averaged across observers (**C.**), across observers and trials of the same type (**D.**), across observers and all trial types (**E.**) The only non significant result is labelled as *n.s.*. All p-values have been calculated via the Wilcoxon rank sum test. No subject was excluded from the analysis. Panels **F.**, **G.**, and **H.** show the box-plots corresponding respectively to panels C., D., E. The green points label the outliers.

Old and young participants map grades were also significantly different. Three examples are shown in Fig. 4. When averaging maps scores across graders, we found a mean score of 17*/*20 ± 0.1 for young, versus 5*/*20 ± 1 for old subjects (Wilcoxon rank sum test, *p <* 10^*-*4^), indicating a much higher accuracy in the maze rendering of the young population. The standard deviation of the scores indicates that the evaluation of old subjects maps was, as expected, much noisier.

Finally, we checked and found that the three performance measurements were significantly correlated. (Spearman test; correlation Δ*d*_*ijk*_-pointing: *p <* 10^*-*6^; correlation Δ*d*_*ijk*_-map: *p* = 0.005; correlation pointing-map: *p* = 0.002; note that the correlation is calculated using Δ*d*_*i*1*B*_, i.e. only the first repetition of trial *B*).

Taken together, theses results consistently point to a significant and detrimental effect of age on the ability to build up a coherent representation of the environment.

### Eye-movements results

Having conducted the experiment in a virtual simulator provides the advantage of isolating the visual contribution to the navigation task; in the absence of proprioceptive and vestibular signals, observers could rely exclusively on their sight to collect information about the environment. Given that, we expected to find a manifestation of learning and/or of the ongoing decision making process in the dynamics of the observers’ eye movements.

In particular, we anticipated significant changes in gaze direction distributions when comparing the exploration phase to the test trials within the same population. During exploration we expected gaze to be more dispersed, with participants trying to maximize the range of their eye movements to probe the unknown environment. During test trials, instead, the uncertainty should be reduced since participants had gained some insight about the environment, had a precise task to accomplish and were not expected to visually explore the environment anymore. Consequently, gaze directions distribution during test trials should be more peaked.

In accordance with our hypothesis, by comparing the gaze distibution’s mean, standard deviation and kurtosis in each sliding window of the region of interest (see Procedure), we found that the gaze direction distributions of the young population was flatter during exploration and more picked during test trials. Interestingly, the opposite was observed for the elderly population (a three way ANOVA was performed with three factors: age (old/young), correctness (correct/incorrect), epoch (exploration/tests). Significant differences between exploration and trials were found for the standard deviation and the kurtosis of horizontal gaze distributions for young vs old participants; results not shown). A possible interpretation of these results could be that young participants were more exploratory at first (flat gaze distribution), while they rather exploited once learning had taken place (picked gaze distribution). To the contrary, old people appeared to be less exploratory in the beginning (picked gaze distribution). This behaviour could have led to some sort of confusion during test trials and, consequently, to a flatter gaze distribution. It is worth mentioning that, while these effects were significant for age, they were not when considering correctness as a factor. Subsequently, we analyzed gaze direction distributions of young and old populations virtually walking trough the maze region of interest while performing the test trials. In agreement with the spontaneous anticipation of locomotor trajectory by gaze direction (Bernardin et al. 2012), we found that also virtual trajectories, or, in other words, joystick turns, were anticipated by gaze direction in both populations. More importantly, we hypothesised that such a gaze anticipatory effect could be delayed in older adults, as a manifestation of a lingering onset of the decision making process. Three principal gaze directional patterns were expected close to the decision point: left, forward and right oriented gaze directions, corresponding to the three possible paths to the goal. However, if we choose to analyze, for example, *B* trials alone, we should expect a bias of gaze direction towards the right, at least for the population who succeeded in performing the task (i.e. who chose more often the right path). While still considering young and old data categories separately, we analyzed two datasets of gaze directions, pooling together *B* trials repetitions: ‘All’ and ‘Correct’ data. ‘All’ data refers to the use of gaze directions drawn from all subjects in the considered category, disregarding whether they succeed or not the trial, whereas ‘Correct’ data refers to the selection of successfully completed trials.

In order to evaluate the onset of the gaze anticipatory effect in the two populations, we evaluated the location within the region of interest 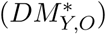 at which the gaze direction distribution significantly departed from unimodality (Hartigan test). As a matter of fact, as soon as subjects got closer to the *FDP*, their gaze direction was no longer peaked at 0°, i.e. towards the central corridor, but started to shift towards the right or the left, i.e. towards the two only available paths in *B* trials, depending on which direction of motion was subsequently chosen. When considering All data populations, we found, as expected, that 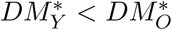, suggesting that young observers were faster in the decision making process. Note that we constrained the speed of the virtual navigation, which was constant and the same for all subjects, hence the result did not depend on a faster motor response in young adults. As evidence that the delayed onset of gaze anticipation was not due to the uncertainty associated to unsuccessful trials but was rather a true age effect, we found that 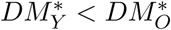 even when we exclusively considered successful trials (see Fig.12 in Appendix).

Our main hypothesis on eye movements pertained to their statistics, rather than to their dynamics. In particular, we expected the frequency of gaze directions towards a given path to be a proxy of the amount of reward attributed to it; in other words, we hypothesised that while planning an alternative route, observers look more often in the direction from which they expect to attain the highest reward, which, in our case, coincides with the direction that they believe leads to the Goal faster. To test our hypothesis, we included in a model-based hiererchical *RL* algorithm (DQ-HRL) the updating of the value of the states observers looked at while moving towards the *FDP*; we assumed that those are the states observers are relying on to mentally plan their detour to the Goal. In the following we describe in detail how the model works.

### Model Results

We first tested the ability of HRL to learn the shortest -and hence most rewarding-path to the Goal in the Tolman maze. As expected, we found that DQ-HRL learned faster and more accurately than both MF-HRL and a standard MB algorithm, which used single-step actions instead of options (i.e. Dyna-Q; Fig. 7B). In DQ-HRL, *Q* values associated to the first decision point attained their optimal values already after the completion of 15 episodes (Fig. 7C., right panel). *Q* values in Dyna-Q also reached their asymptote quite fast, i.e. starting from 20 episodes, however, the algorithm was less accurate than HRL, since it could not disentangle most actions values (i.e. *Q*(*L*) = *Q*(*D*) = *Q*(*R*) in Fig. 7C., left panel). The model free version of HRL was the slowest and less accurate (Fig. 7C., central panel).

**Figure 7:**
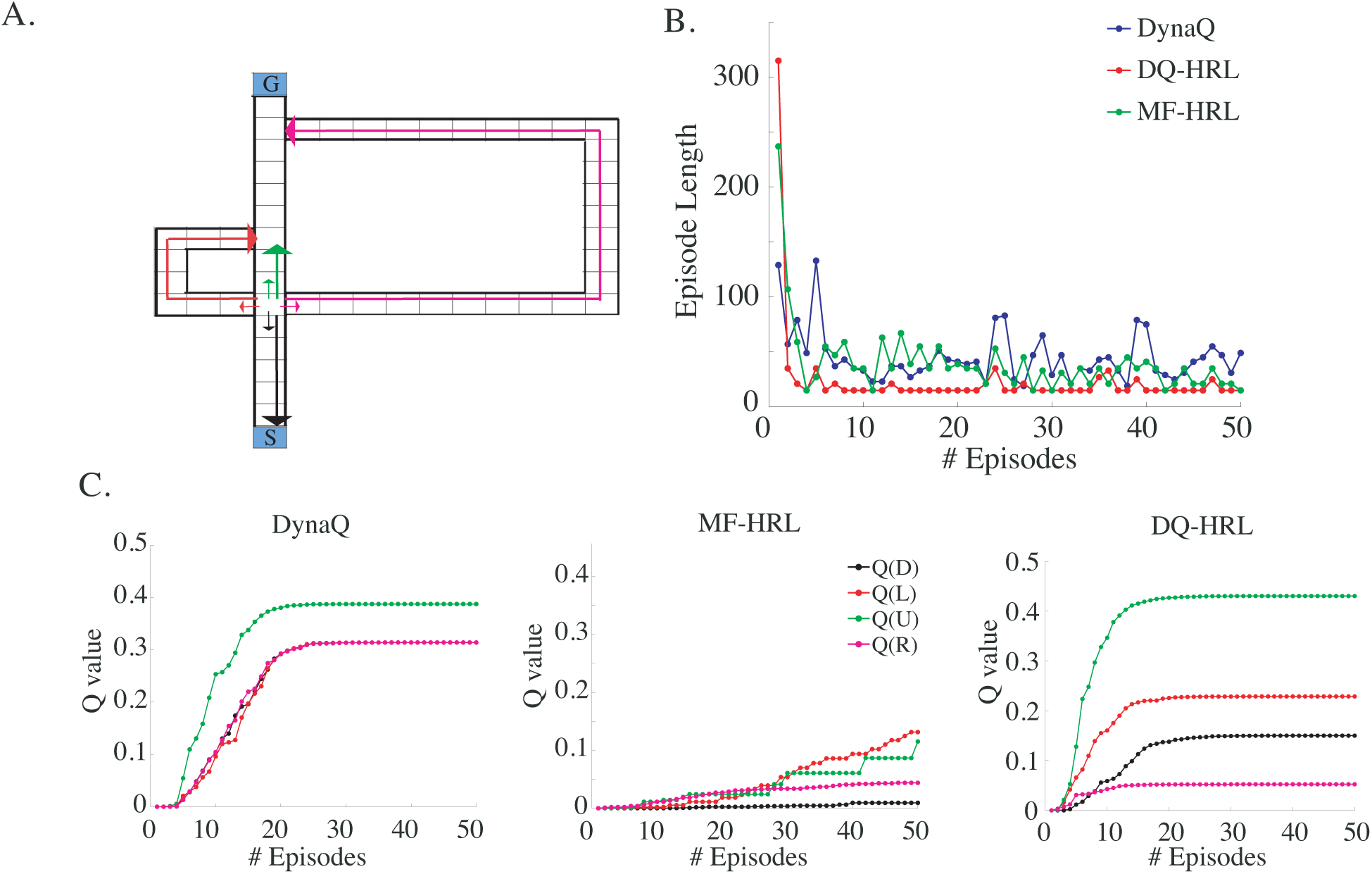
A. Cartoon of a subset of available *options*, used in *HRL* algorithms, versus *primitive actions*, used in standard *RL* algorithms. The former are a spatially extended version of the latter. **B.** Performance of model based *HRL*, model-free *HRL* and model-based standard *RL* (i.e. Dyna-Q) measured as in Eq. 1 for the Tolman maze shown in A. **C** Dynamics of *Q* values in *FDP* for the three algorithms shown in B. The optimal algorithm should converge to *Q*(*U*) *> Q*(*L*) *> Q*(*D*) *> Q*(*R*), as *DQ - HRL* does. Parameters: *α* = 0.1; *γ* = 0.9; *τ* = 0.1; *n*_*pl*_ = 10 for model based; *n*_*pl*_ = 0 for model free.

Importantly, the way in which we defined options in HRL algorithms impedes the agent to repeatedly go back and forth in small subregions of the maze, i.e. to get jammed in unrealistic loops, which we never observed in the subjects behaviour. Those unrealistic loops are hardly avoidable in standard MB algorithms, which use primitive actions.

Subsequently, we used DQ-HRL in a two-steps procedure to reproduce the performance of young (*Y*) and old (*O*) subjects during the execution of the test trials (Fig. 6B). First, we fed the model with the actual sequence of states and options taken by each subject during the *exploration* phase. Initial *Q* values were set to zero for all subjects; the values of *α* and *n*_*pl*_ were fixed and assumed to be equal for all subjects within a population; the noise parameter *τ* was unnecessary since state transitions in the model were not governed by any stochastic policy, but were deterministically defined by the actual trajectory followed by the subjects. The reward *r* associated with the very last transition bringing the agent to the Goal was set to one; all the other rewards were set to zero.

**Figure 6:**
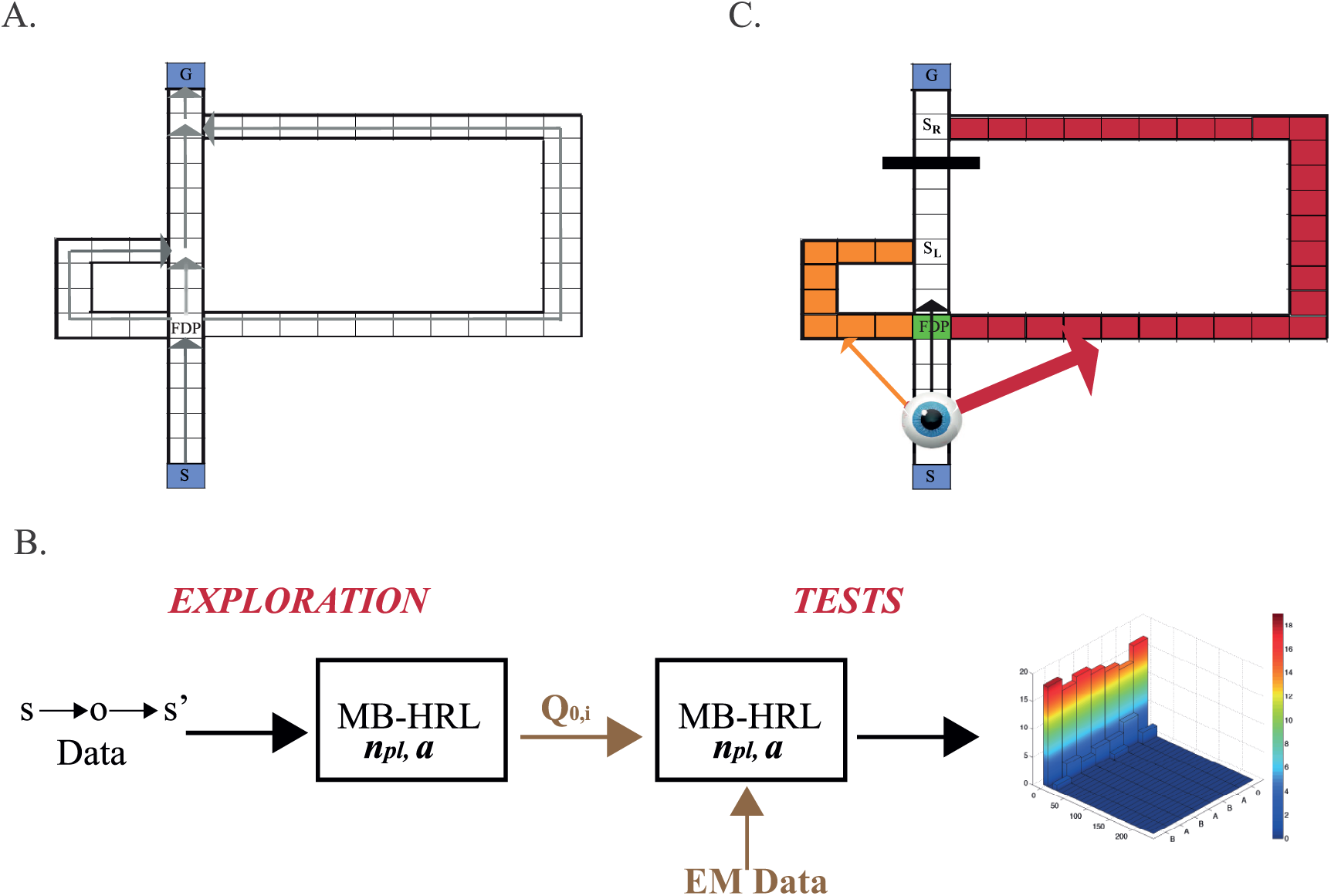
A. For modelling purposes, the Tolman maze is decomposed in a set of 52 states (black squares in the cartoon) and 6 non-overlapping *options* (grey arrows). Each option is bidirectional, although, for clarity, only one direction is depicted in the figure. The length *k* of an option is given by the number of underlying states it covers. For example, the first option initiates at the starting point of the maze (*S*) and terminates at the first decision point (*FDP*), hence *k* = 6. **B.** The model was first “fed” with the individual subjects trajectories measured during the exploration phase, and produced the *Q* values of each individual subject as output. Those values, together with individual subjects eye-movements, were used to probe the model with the sequence of behavioural test trials 0-A-B-A-B-A-B. In both the exploration and the test phase the relevant parameters were the number of planning steps *n*_*pl*_ and the learning rate *α*. **C.** Subjects eye-movements were used in the model to modulate the reward associated with the options available at *FDP* during the test trials. In the cartoon a *B* trial is depicted. If a subject, navigating from *S* to *FDP,* fixated 20% of the time towards the right, the reward associated to the transition *FDP* → *s*_*R*_ was set to *r* = 0.2. Likewise, if the proportion of fixation to the left was 10%, the reward for the transition *FDP* → *s*_*L*_ was set to *r* = 0.1.

The feeding procedure provided, for each subject, the *Q* values at the end of the exploration phase, i.e. *Q*_0*,i*_ (Fig. 6B).

The following step of the procedure consisted in *i)* setting individual *Q*_0*,i*_ as initial conditions for running the model with individual *test trials*, and *ii)* using subjects gaze directions during the execution of test trials as a tool to re-plan a path within the maze. The parameters *α* and *n*_*pl*_ were assumed to remain unchanged for both populations with respect to the exploration phase; furthermore, we set *τ ≃* 0 since, contrary to the exploration phase, during the tests trials we asked subjects to be “greedy”, i.e. to reach the Goal via the shortest possible path.

We run the model for each individual subject in the seven test conditions 0 *-A-B-A-B-A-B*.

To clarify the role of eye movements (*EM*) in the model, let us describe the procedure we followed for a given subject *i* engaged, for example, in a *B* trial. *Q* values were initialised at *Q*_0*,i*_. Starting from *s* = *S*, the agent chose an option according to the policy in Eq.2, where *Q*(*s, o*) = *Q*_0*,i*_(*s, o*), and landed in *s’* = *FDP,* i.e. in the maze first decision point. The value of *Q*_0*,i*_(*s, o*) was updated according to Eq. 3 and the state transition was stored in *M* together with its associated reward *r*:

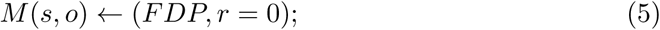

we assumed that, before choosing any other option, the agent used subject *i* eye movements for planning his next move. From the eye-tracking data, we extracted the number and the direction of fixations made by subject *i* while navigating from *S* to *FDP* (Fig. 5). If, for example, 70% of the fixations were made straight ahead towards the corridor, 20% towards the right arm and, 10% towards the left arm, we stored in *M* the following transitions:

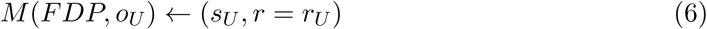

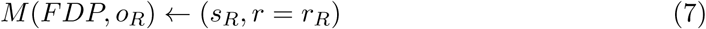

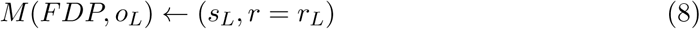

where *o*_*U*_ labelled the option straight ahead (or *UP*) and analogously, *o*_*R*_ (*o*_*L*_) the option right (left). The state *s*_*U*_ labelled the terminal state of *o*_*U*_ coming from *FDP* (analogously for *s*_*R*_ and *s*_*L*_; Fig. 6C). Importantly, the value of the reward associated to each transition was set equal to the proportion of EM made in the corresponding direction; in our example, *r*_*U*_ = *-*0.7, *r*_*R*_ = 0.2 and *r*_*L*_ = 0.1. Note that we set *r*_*U*_negative for all trials in which a block was used (i.e. *A* and *B* trials); we reasoned that, since the block was made clearly visible, subjects were well aware of the impossibility to choose *o*_*U*_ in those cases (at the expenses of bumping straight into the block) and internally associated it with a negative reward.

**Figure 5:**
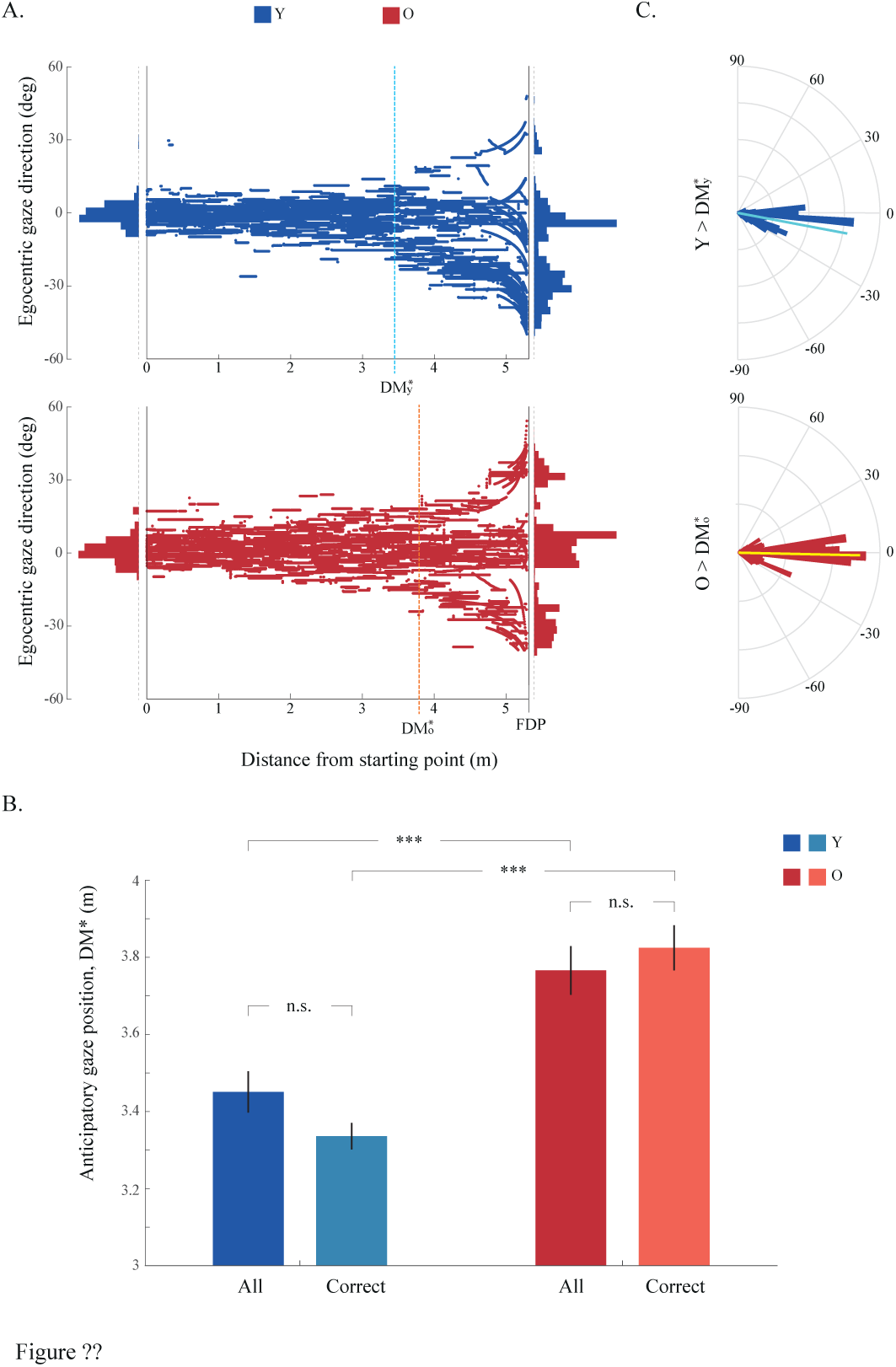
A. On the left plot, gaze direction distributions of ‘All’ Y and ‘All’ O data binned over a sliding window as a function of the distance between *S* and *FDP.* 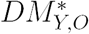 labels the location at which the distribution switches from unimodal to multimodal, i.e. the location at which the eyes anticipated the direction of motion. On the right plot, the gaze direction distribution obtained in the last bin, i.e. closest to the *FDP,* is represented on a semi-polar plot; the superimposed line labels the median of the distribution. Note how the gaze distribution of Y observers is much more biased towards the right direction. **B.** To estimate the variability associated to the decision making point 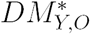, we applied for each of the four datasets Young correct (YC)/old correct (OC), Young All data (YAD) / old All data (OAD) the following procedure. For each sliding window, we generated a kernel probability density estimate for the current gaze direction distribution, using a normal kernel smoothing function. We then randomly sampled 20 synthetic datasets from the pdf of the kernel distribution. For each dataset, we performed the Hartigan test. Parametric t-tests showed that ‘All’ Y and ‘All’ O data exhibited a significant (*p <* 0.001) difference between onsets of the multimodal gaze direction pattern. Within each of these groups, changes in onset value when considering ‘All’ or only ‘Correct’ trials were not significant.

While still in *FDP,* the agent randomly sampled *n*_*pl*_ states and options from *M* and updated the corresponding *Q* values. Crucially, in our procedure EM allow the agent to do planning using a larger sample of transitions (i.e. those associated to visited states, as well as those associated to states which were only looked at) and, eventually, end up with a better and faster estimation of *Q* values; without EM, instead, the agent would rely only on a single transition (Eq. 5) for planning while in *FDP.*

When the *n*_*pl*_ number of planning steps were over, the agent chose almost always (since *τ ≃* 0) the option associated with the highest *Q* value, *o*_*R*_ in our example, ending up in *s*_*R*_. Once the agent went past *FDP,* we reasoned that EM would no longer reflect a planning strategy to reach the Goal and hence we did not consider subjects EM (i.e. beyond *FDP,* only those transitions which were actually executed were stored in *M* and used for planning, as in standard MB frameworks). The test trial ended as soon as the agent reached the Goal state. The model performance was quantified as in Eq. 1 and compared with the behavioral data: consistently with the data, we plotted the histogram of Δ*d*_*ijk*_ for both populations and found that the model behaved qualitatively very similarly to the subjects (Fig. 8A, for a given set of parameters). Box-plots, showing Δ*d*_*ijk*_ medians across observers and trials, confirmed this result also quantitatively (see inset in Fig. 8B).

**Figure 8:**
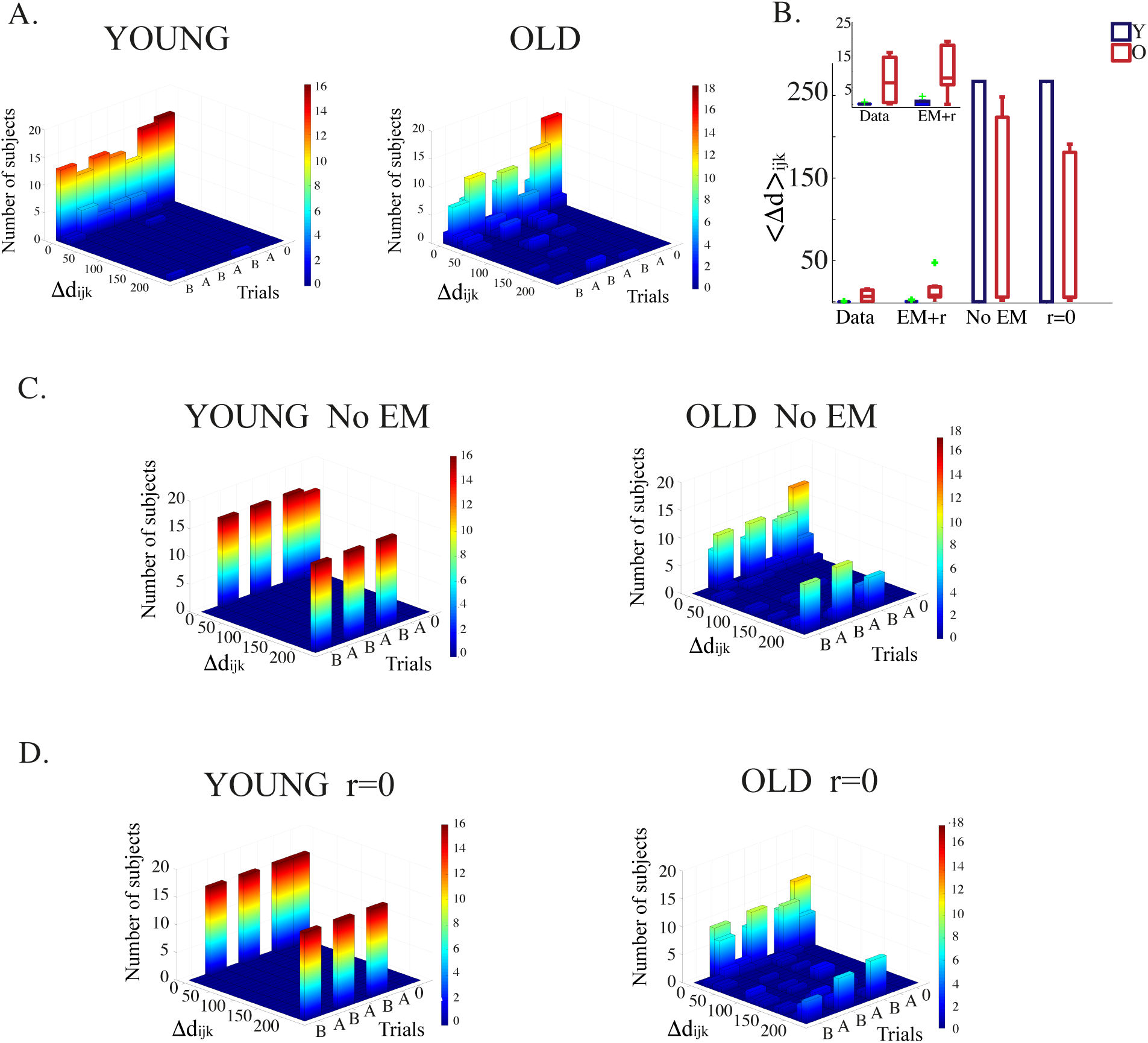
A. Model performance measured, analogously to the data, as the distance travelled by individual observers in each test trial. To obtain these results, individual exploration data as well as individual EM have been injected in the *HRL* algorithm. **B.** Median across observers and trials of the travelled distance, calculated for the data, for the model using EM statistics and associated reward modulation, for the model without EM and for the model in which EM do not entail a reward modulation. Disregarding EM and/or reward modulation is significantly detrimental for the model performance. **C.** Same as in A., but for the model which does not use EM. **D.** Same as in A., but for the model which use EM only to select paths to plan upon, without associating any reward to them. Parameters: young population: *α* = 0.21; *γ* = 0.9; *τ* = 0.01; *n*_*pl*_ = 29; old population: *α* = 0.01; *γ* = 0.9; *τ* = 0.01; *n*_*pl*_ = 17;

Importantly, when EM were discarded, the model was unable to reproduce observers performance with the same accuracy; in the case of young subjects, while the algorithm performed optimally in 0 and *A* trials, its performance was severely suboptimal in *B* trials (left panel in Fig. 8C). The reason is fairly straightforward: at the end of the feeding procedure (i.e. at the end of the exploration phase), *Q* values of individual young subjects had, on average, converged to their optimal value. For the values associated with *FDP,* for example, *Q*(*U*) *> Q*(*L*) *> Q*(*D*) *> Q*(*R*) (Fig. 9C). Accordingly, in the upcoming trial 0, the agent most likely chose option *U* while in *FDP* (since *τ ∼* 0), and successfully reached the Goal travelling trough the central corridor. In *A* and *B* trials, instead, *U* was negatively rewarded because of the block, and the agent chose the *L* option as the most rewarding one. The *L* choice successfully led to the Goal in *A*, but not in *B* trials. To correctly perform in B trials, standard RL algorithms should re-learn the new configuration of the environment, once blocks are introduced. The same reasoning holds true for old subjects; in this case, however, the variability of the algorithm performance was higher, given that *Q* values did not yet converge to their optimal values at the end of the exploration phase (right panel in Fig. 8C).

**Figure 9:**
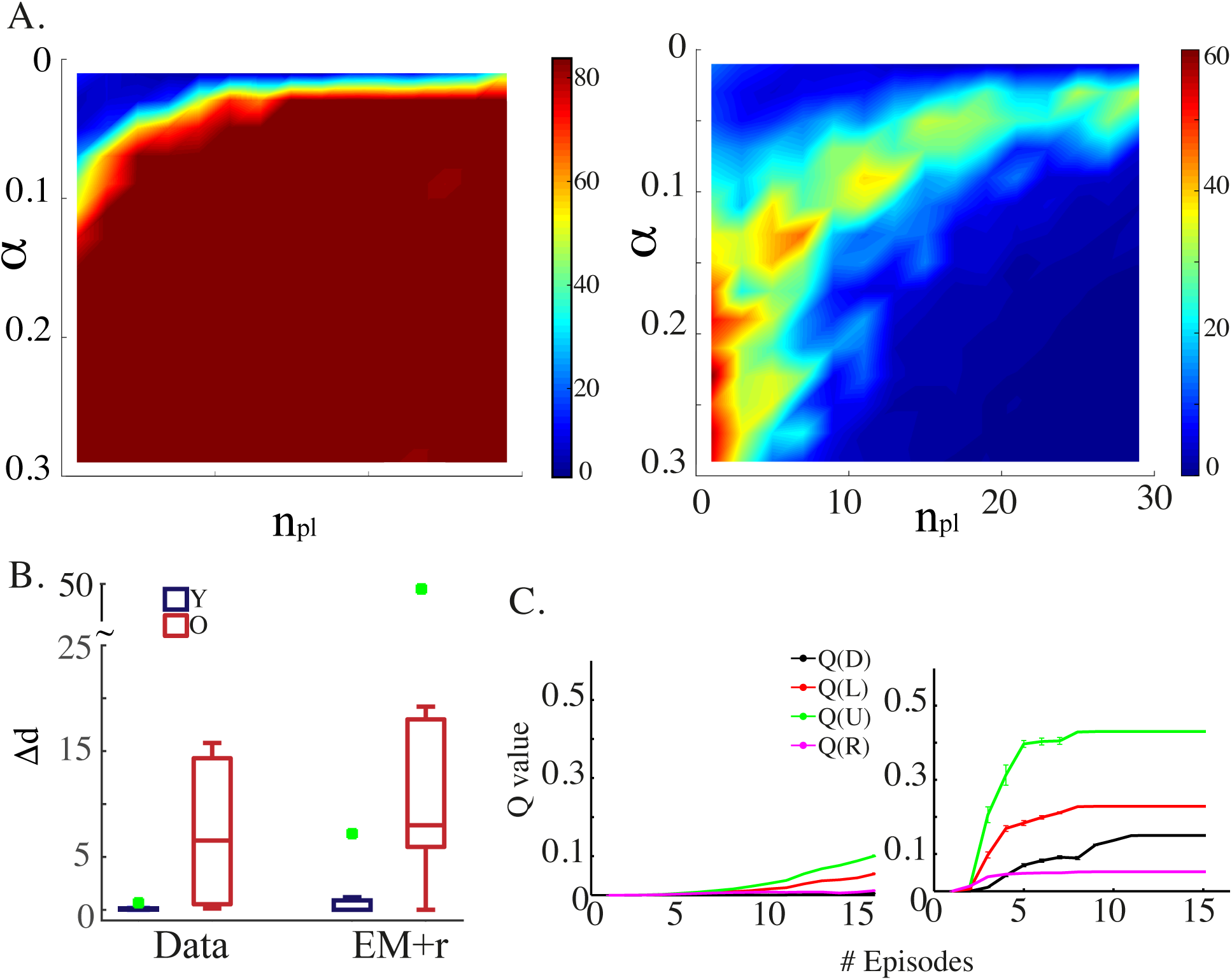
Sensitivity anlaysis A. Difference in performance between data and model, calculated as in Eq. 12, as a function of the learning rate *α* and the number of planning steps *n*_*pl*_. The best parameters are: young population: *α*^∗^ = 0.19;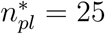; old population: *α*^∗^ = 0.01; 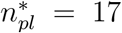. **B.** Median across observers and trials of the travelled distance, calculated for the data and for the model with *α* = *α*^∗^ and 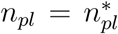. Average *Q* values dynamics in *FDP* as a result of the feeding procedure in which we set *α* = *α*^∗^ and 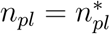. Error bars represent s.e.m. For the exploration data used in the feeding procedure, the average number of episodes across observers was 10.27+2.53, while the average number of state transitions (i.e. the average number of options taken) was 64.16+12.21.

A similar suboptimal behaviour was found when EM were used to identify available paths to plan upon, but not to assign their corresponding rewards (Fig. 8D). In this case, Eq. 8 become:

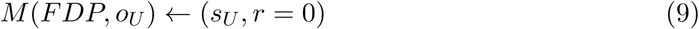

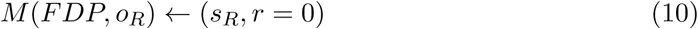

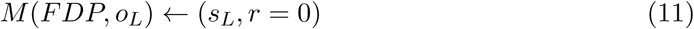

and, as before, the environment should be re-learned for the algorithm to successfully perform on *B* trials.

Up to know we showed the model results for a single set of parameters. Next, we studied in which region of parameters these results hold. The relevant parameters were *α*, i.e. the learning rate, and *n*_*pl*_, i.e. the number of planning steps. To compare the model performance to the data we considered the following quantity, for the young and the old population separately:

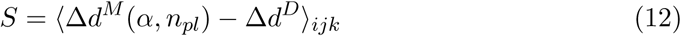

where *k* labelled the trial type (*k* = 0*, A, B*), *j* the number of repetitions of each *k* trial (*j* = 1, 2, 3), *i* the subject; the superscripts *M* and *D* indicated, respectively, model and data. In other words, we calculated the median of the deviation from the minimum path over subjects and trials for any given pair of parameters (*α, n*_*pl*_) and compared it to the data; accordingly, the best parameters were those which minimized *S*.

We found that, young subjects data were best described by both large *α* and large *n*_*pl*_; old subjects data instead were best described by smaller values of both parameters (blue regions in Fig. 9A). Consistently, plotting the model performance, 〈Δ*d*^*M*^ 〉 _*ijk*_, for the best parameters yielded results which were very close to the data for both populations (Fig. 9B). Note that, for the young population, the 0 *< α <* 0.1 region in the parameter space for which *S* appeared to be small is actually a local minimum (see Fig. 10 in Appendix).

**Figure 10:**
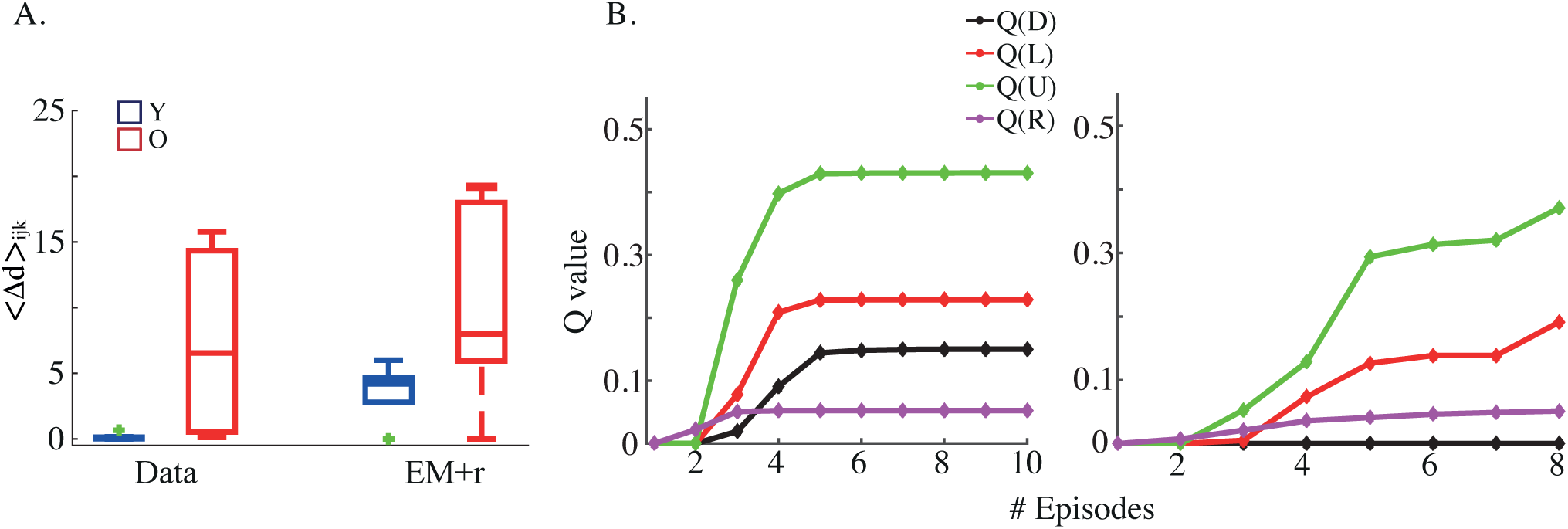
A. Median across observers and trials of the travelled distance, calculated for the data and for the model with *α* and *n*_*pl*_ set at the local minimum for the young population, which corresponds to *α* = 0.01; *n*_*pl*_ = 15; **B.** Left: Average *Q* values dynamics in *FDP* for the best old subject, for whom *α* = 0.21 and *n*_*pl*_ = 21. Right: Average *Q* values dynamics in *FDP* for the worst young subject, for whom *α* = 0.21 and *n*_*pl*_ = 3.

We then looked at the dynamics of Q values in *FDP* obtained from the feeding procedure; we found that the parameters which best described the data were also those for which the *Q* values of the young population were optimal well before the end of the exploration phase, while those of the old population never reached optimality (Fig. 9C). This partly accounts for the flawless performance of the model in reproducing young data in the test trials and it provides an indication that young observers learned the structure of the environment already before the end of the exploration phase. Such a result was corroborated by verbal reports of young subjects at the end of the exploration phase, who spontaneously stated that 12min of exploration were unnecessary since they understood the maze structure already after the first couple of minutes.

The ensemble of these results quite naturally leads to the following interpretation: old subjects are slower learners (small *α*) and they plan their way through the maze less extensively (smaller *n*_*pl*_). As a consequence, their behaviour is much closer to that of a model free algorithm rather than model based, as exemplified by their *Q* values dynamics (compare Fig. 9C. with Fig. 7C). It is worth noticing that the performance of the best observer among the elderly was best described by a much bigger *n*_*pl*_ than that describing the performance of the worst young subject. It follows that an optimal subject is one who plans his way through the maze more exhaustively (Fig. 10B in the Appendix).

## Discussion

Despite the vast literature on spatial navigation, not much is known about how the brain adapts to unexpected changes in the environment (for a recent review see Spiers and Gilbert 2015); even less is known about how this ability is effected by a non-pathological aging process. Here, we relied on a RL framework to provide a mechanistic interpretation of how a detour task is solved. The model quite successfully reproduced the age related differences we found in the human ability to solve a Tolman detour task, in particular the impediment of older observers to grasp which alternative route would bring them faster to the Goal. The model supports two pivotal assertions on the role of eye movements in human planning strategies: i) the pattern of eye movements reflects the memory replay of experienced paths, which serves the planning of alternative routes; ii) the statistics of eye movements, i.e. the frequency with which observers look towards a given path, is a measure of the reward they expected when following it. The model’s parameters which best fit the data indicated that older adults are slower learners but, indeed, they plan alternative paths through the maze like younger ones, although to a slightly less extent.

In the light of these results we put forward that global planning strategies do not radically change with age and that the impediment in solving a detour task found in older adults might rather be due to the building up of the wrong internal representation of the environment. In other words, older adults do plan, but over a wrong cognitive map.

Although it remains speculative, this hypothesis gets support from the maze sketched maps drawn by our older subjects, as well as from evidence that it takes a longer time for older adults to create an internal representation of the environment while navigating (Iaria et al. 2009). Consistently, the time we allocated for exploration in our paradigm might have been insufficient for older observers to build up a correct map of the maze. Moreover, older participants’ pattern of eye movements points to a suboptimal strategy of acquisition of information about the environment during the exploration phase, which could also contribute to a defected map formation.

Crucially, we checked that the inability we found in older observers was not exclusively due to the lack of both proprioceptive and vestibular signals, which typically provide a reliable support to spatial navigation; when we asked observers to solve the Tolman detour task while actually walking through it, we found just about the same results obtained in the virtual setting (results not yet published).

### On eye movements

An abundance of studies have shown that older adults are significantly impeded in spatial navigation tasks (for reviews see Moffat 2009; Lester et al. 2017). A bunch of these contributions measured performance differences between younger and older adults in tasks which supposedly relied on the ability to use and/or to form a cognitive map, finding, as we did, a detrimental effect of aging on spatial navigation performance (Newman and Kaszniak 2000; Moffat and Resnick 2002; Iaria et al. 2009). There is, however, to the best of our knowledge, no study which measured performance in a Tolman detour task in both younger and older observers and, more importantly, which analyzed the role of eye movements during a route re-planning strategy.

In the literature, eye movements have been strongly implicated in memory encoding and retrieval; for example, limiting the number of fixations appears to be detrimental for both, when compared to free-viewing conditions (Henderson et al. 2005; Johansson and Johansson 2014). Moreover, altered fixation patterns in older adults have been proposed to be functionally linked to memory deficits (Chan et al. 2011; Hannula et al. 2007; Olsen et al. 2015; Ryan et al. 2000; Rondina et al. 2017; Shih et al. 2012; Voss et al. 2011). Consistently, we found that gaze direction distributions differed significantly between young and old observers during both the exploration and the probe trials (although we did not find any difference in the sheer number of fixations made; results not shown). In particular, the pattern of fixation of older observers exploring the maze, points to an inefficient encoding of stable information about the environment; likewise, the pattern of fixations recorded during the execution of the test trials is consistent with a suboptimal retrieval process, which, in older adults, leads up to a delayed decision about which path to choose (Fig. 5).

If the link between eye movements and memory is fairly well documented, much less is known about eye movements in the context of the re-planning of alternative routes. Two important pieces of evidence related eye movements and activity in the hippocampus, one of the brain structure most strongly implicated in the replay of past experiences as well as in the planning of future events (Buckner 2010; Foster and Wilson 2006; Gelbard-Sagiv et al. 2008; Javadi et al. 2017; Johnson and Redish 2007; Pastalkova et al. 2008; Pfeiffer and Foster 2013; Schacter et al. 2012; Wu and Foster 2014). First, during the encoding of novel stimuli, the number of fixations correlate with hippocampus activity, suggesting that visual sampling might be directly related to the formation of representations in the hippocampus (Liu et al. 2017). Second, the hippocampus guides where to look during memory retrieval (Hannula et al. 2007; Hannula and Greene 2012; Ryals et al. 2015; Ryan et al. 2000). These findings appear to be fully consistent with what we propose: eye movements subserve the retrieval of experienced paths from a mental representation of the environment, upon which future routes are planned. In other words, eye movements are the explicit manifestation of hippocampus-based computations implicated in planning. Our theory pushes the link between eye movements and planning even one step further: gaze directions associated with planning reflect the expectation of future rewards, meaning that we plan more often upon routes which we believe to be more rewarding and which, consequently, we deem more valuable. Furthermore, solving the detour task imply a re-evaluation of the value of the options available after the alteration of the environment (i.e. the introduction of the block in the maze); such an updating of reward assignment has been proposed to implicate hippocampal-striatal connectivity (Pennartz et al. 2011; Wimmer and Shohamy 2012).

Note that, here, we implied that planning consists in two concurrent processes i) the memory replay of experienced paths; ii) the re-evaluation, crucial for future decisions, of the benefits associated with any of these paths. Nonetheless, we acknowledge that planning does not necessarily rely on the recall of past experiences, e.g. in some circumstances, hippocampal cells are able to generate firing sequences representing paths never taken by the animal (Gupta et al. 2010; Ó lafsdóttir et al. 2015). Moreover, it is important to stress that planning, in particular in the context of detours, is supported not only by the hippocampus but also by prefrontal cortices, although the relative contribution of the two is still unclear (Martinet et al. 2011; Spiers and Maguire 2006; Spiers and Gilbert 2015; Javadi et al. 2017). Furthermore, both ventromedial prefrontal and striatal areas have been suggested to code for the value of the reward associated to mentally planned scenarios (Benoit et al. 2014; Lin et al. 2015).

### On modelling

Reinforcement learning has been widely used as a learning rule in spatial navigation, not only for its efficiency, but also on the account of its biological plausibility. Mid-brain dopaminergic neurons in the substantia nigra pars compacta (SNc) and ventral tegmental area (VTA) supply the striatum with dopamine, signalling a reward prediction error (Schultz et al. 1997), which represents the core of temporal-difference (TD) RL algorithms. The striatum itself is thought to be responsible for the computation of stimulus-response associations typical of MF versions of RL (Gläscher et al. 2010; Johnson et al. 2007; van Der Meer and Redish 2011). The more flexible learning rule implemented by MB algorithms, which relies on an explicit internal model of the environment, is instead carried out by the hippocampus (Gustafson and Daw 2011; Hasselmo 2005; Hirel et al. 2013; Martinet et al. 2011; Simon and Daw 2011).

Recently, the literature has converged on the idea that a combination of both MB and MF algorithms is most suitable to describe human navigation strategies, or, more generally human sequential decision making (Daw et al. 2005; Daw et al. 2011; Wunderlich et al. 2012; Gershman et al. 2014; Lee et al. 2014; Otto et al. 2014; Lee and Keramati 2017). In particular, the Dyna-Q algorithm (Sutton 1990) has been shown to be an optimal candidate, given that it relies by construction on a mixture of the two: the MB system, using an internal model of the environment, simulates offline state action transitions whose *Q* values are then updated through the MF system via a TD algorithm (Gershman et al. 2014).

**Figure 12:**
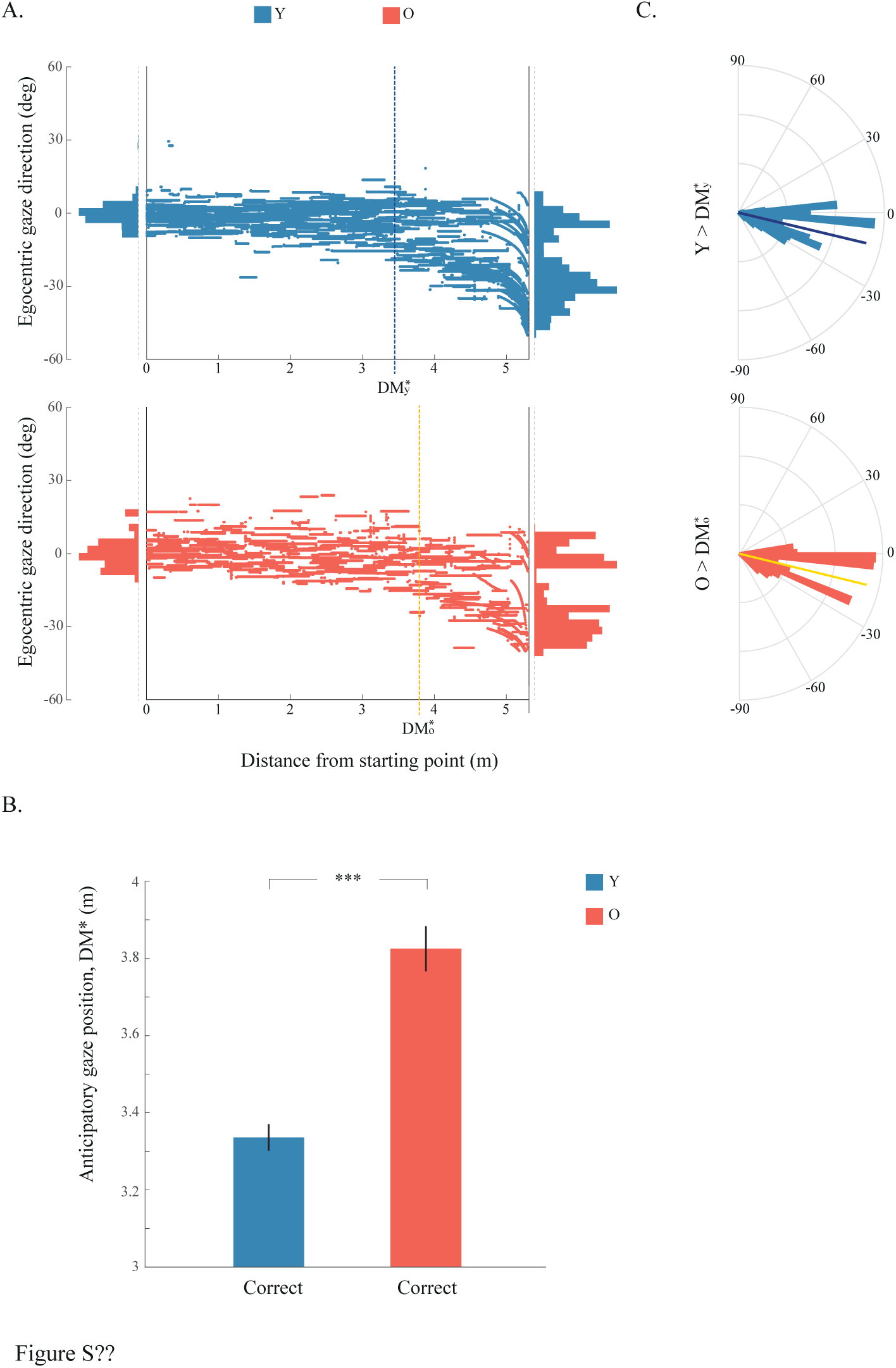
Same as Fig. 5 but for ‘Correct’ Y and ‘Correct’ O datasets.

**Figure 11:**
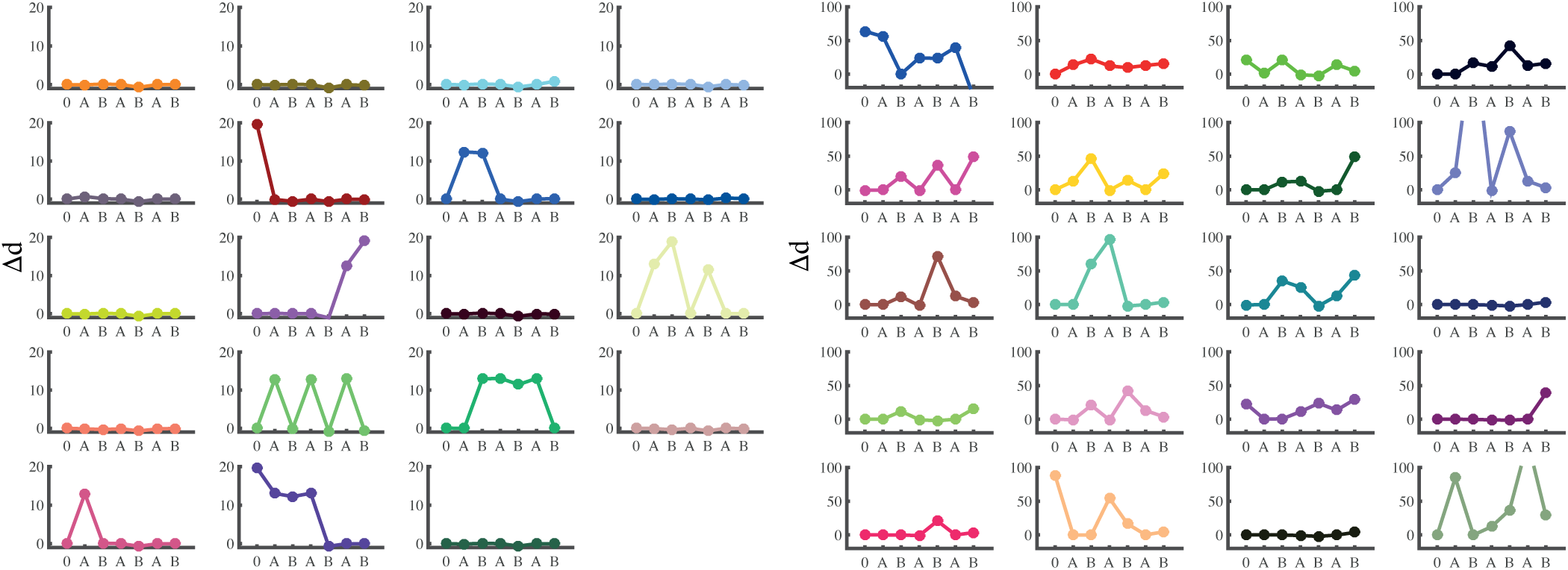
A. B..

Here, seeking computational efficiency, we implemented Dyna-Q in the framework of HRL (DQ-HRL). The advantage of partitioning the environment- as well as the actions therein-in larger chunks is well known computationally (Badre et al. 2010; Botvinick et al. 2009; Sutton and Barto 1998) and it has recently been corroborated, behaviourally, by the finding that the elaboration of plans in human observers is indeed hierarchical (Balaguer et al. 2016). We showed that DQ-HRL is faster in learning the shortest path to the Goal in a (block-free) Tolman maze, than both classical Dyna-Q and the model free version of HRL (MF-HRL; Fig. 7). This is mainly because classical algorithms need to learn the value of each state and action within each arm of the maze, which would entail wasting time in going back and forth through sub-portions of a selected arm. To the contrary, an algorithm like HRL would learn the value of an arm as a whole, resulting in much faster learning.

However, although extremely efficient, even HRL would need to re-learn the new structure of the environment once the blocks are introduced and, consequently, it would always perform poorly with respect to a young observer, who is capable of taking the optimal detour without further training. In other words, in any RL algorithm the agent would need to visit several time the state immediately close to the block to learn to attribute the appropriate value to it and to be able to choose a suitable alternative (see for example Russek et al. 2017). However, none of our subjects during the test trials needed to do so, i.e. none of our subjects ever walked straight to the block: looking at the obstacle from a distance was enough to grasp that the central corridor was occluded and that one of the other available paths had to be chosen instead. Inspired by this observation, we gave the agent in the DQ-HRL algorithm the ability to plan over the states our observers only looked at, ending up with a RL model which, for the first time, can learn the Tolman detour task in one-shot.

Upon fitting the subjects’ trajectories and eye-movements, our model predicts that older adults are not only slower learners, but also less efficient planners compared to younger adults. Plausibly, ageing has an effect on the quality of the internal representation of the environment on which planning is based. Future work might focus on pushing the eye-movements analysis to a higher level of sophistication, e.g. to understand in which respect a cognitive map might result inaccurate and whether ageing affects specifically the binding of the different structural elements of the environment together or rather other aspects of the map formation.

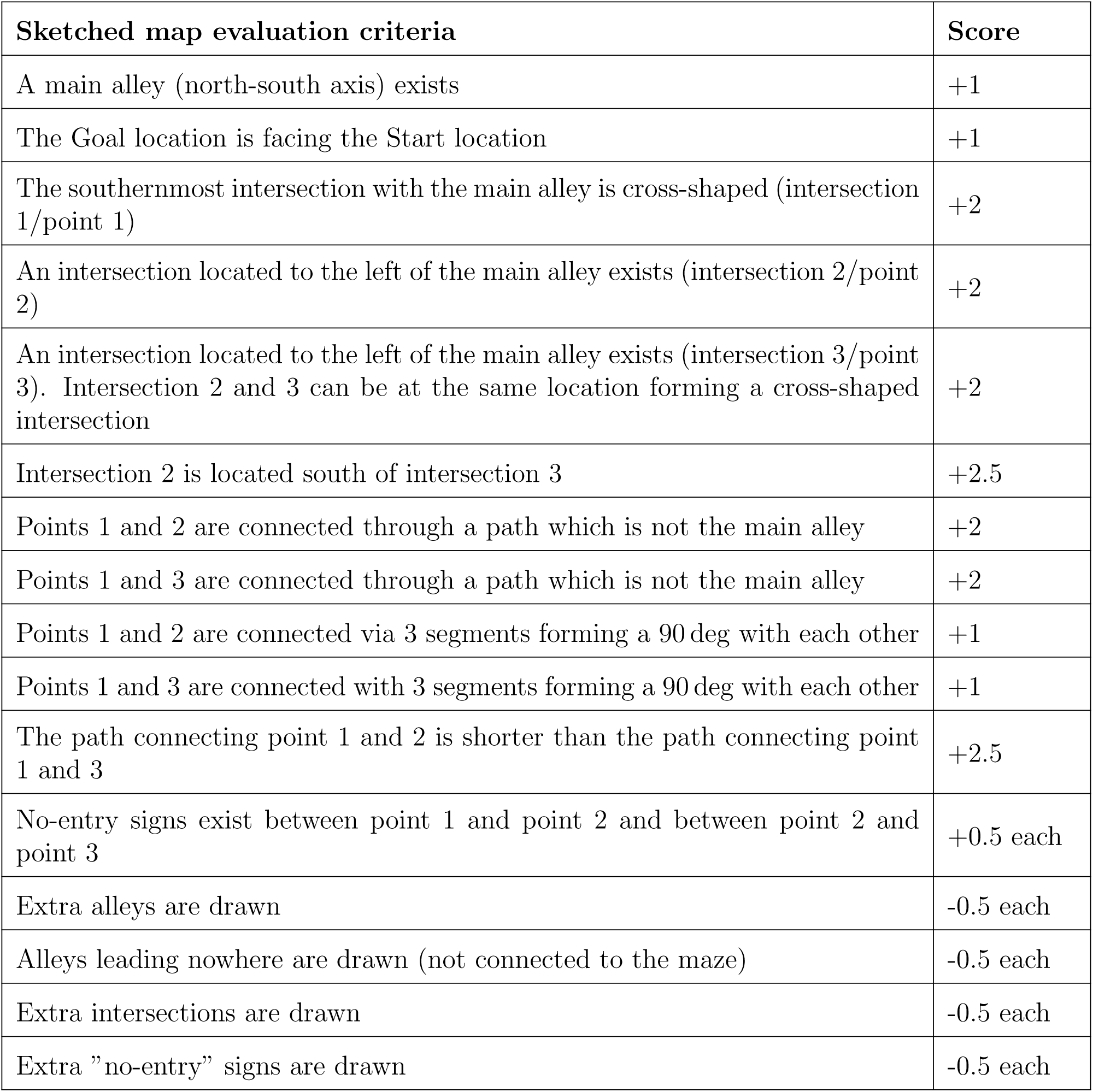

